# On quality thresholds for the clustering of molecular structures

**DOI:** 10.1101/2022.08.31.505983

**Authors:** Xavier Daura, Oscar Conchillo-Solé

**Affiliations:** Catalan Institution for Research and Advanced Studies (ICREA), Barcelona, Spain; Institute of Biotechnology and Biomedicine, Universitat Autònoma de Barcelona, Spain; Biomedical Research Networking Center in Bioengineering, Biomaterials and Nanomedicine (CIBER-BBN), Barcelona, Spain; Department of Genetics and Microbiology, Universitat Autònoma de Barcelona, Barcelona, Spain

## Abstract

It has been recently suggested that diametral (so-called quality) similarity thresholds are superior to radial ones for the clustering of molecular three-dimensional structures. ^1^ The argument has been made for two clustering algorithms available in various software packages for the analysis of molecular structures from ensembles generated by computer simulations, attributed to Daura et al. ^2,3^ (radial threshold) and Heyer et al. ^4^ (diametral threshold). Here, we compare these two algorithms using the root-mean-squared-difference between the Cartesian coordinates of selected atoms as pairwise similarity metric. We discuss formally the relation between these two methods and illustrate their behavior with two examples, a set of points in two dimensions and the coordinates of the tau polypeptide along a trajectory extracted from a replica-exchange molecular-dynamics simulation. ^1,5^ We show that the two methods produce equally-sized clusters as long as adequate choices are made for the respective thresholds. The real issue is not whether the threshold is radial or diametral, but how to choose in either case a threshold value that is physically meaningful. We will argue that, when clustering molecular structures with the RMSD as metric, the simplest best guess for a threshold is actually radial in nature.

## Introduction

Over two decades ago, Heyer et al. ^4^ developed an algorithm to cluster open reading frames (ORFs) based on their expression levels, using what they came to call jackknife correlation as pairwise similarity metric. The focus of the algorithm was to find large clusters that had a “quality” guarantee, i.e., a minimum jackknife correlation between any two ORFs belonging to the same cluster. In other words, clusters would be guaranteed to have a maximum diameter, determined by a correlation threshold, ensuring the transitive property for the relation ‘correlation > threshold’ between element pairs in a cluster (i.e., if correlation(*a, b*) > threshold and correlation(*b, c*) > threshold, then correlation(*a, c*) > threshold, for any *a, b, c* belonging to the cluster). The algorithm was named quality cluster (QT_Clust) and it has since been used for the clustering of several other types of data, including molecular structures.^1^ In this study we will refer to this algorithm as QTC.

The same year, Daura et al. ^2,3^ introduced an algorithm to cluster molecular structures using the root-mean-squared-difference (RMSD) between the Cartesian coordinates of selected atoms as pairwise similarity metric. The algorithm was meant to favor the most populated cluster and was radial in nature, i.e., the RMSD threshold was applied from a configuration taken as cluster reference, meaning transitivity was not ensured for the relation ‘RMSD < threshold’. In this study we will refer to this algorithm as RTC. The two algorithms are described under Computational details and analyzed formally under Results and discussion.

When applied to molecular structures using the RMSD as metric, both QTC and RTC scan a precalculated RMSD matrix in search for the molecular configuration with the largest number of neighbors satisfying the threshold, in an iterative process that outputs a new cluster at each step and ends when there are no further configurations to cluster. The difference between the two algorithms lies in the nature of the threshold (diametral or radial) and the procedure to count the neighbors in. Recently, González-Alemán et al. ^1^ pointed out that, because of their similarity, these two algorithms were often confused in various software implementations commonly used in the field, misleading their users. They also evaluated the performance of the two algorithms by analyzing a trajectory of the tau polypeptide extracted from a replica-exchange molecular-dynamics simulation previously published by Shea and Levine.^5^ In doing so, they used the same RMSD value for the thresholds applied in the two algorithms. They concluded that, due to its lack of a quality threshold, the RTC algorithm (referred to as Daura’s algorithm in the paper) tends to cluster unrelated configurations together, and gave examples in which different QTC clusters were found as composing a single RTC cluster.

Clearly, the results observed by González-Alemán et al. had little to do with the quality of the thresholds and much to do with using the same RMSD value for a radial and a diametral threshold. While the relation ‘RMSD < threshold’ is not transitive for the set of configurations conforming a cluster generated by the RTC algorithm, the alternative relation ‘RMSD < diameter’ is. This leads to the following question: can we generate clusters with a predetermined maximum diameter using the RTC algorithm? In other words, can we choose the radial threshold in the RTC algorithm in such a way that the maximum diameter of a cluster will be equal to the threshold we would use with the QTC algorithm? The answer is of course yes, if one would actually wish to do so.

In the following sections we will present formally and analyze in detail the characteristics of the RTC and QTC methods. Although the two methods have been available and heavily used since over two decades, a detailed analysis of their properties has not been published. We will show that it is indeed possible to obtain equally sized clusters with the two algorithms and will argue that this is in fact of little importance because, first, the threshold is an arbitrary quantity that may have different “ideal” values depending on the objective of the analysis and, second, for the purposes discussed here the simplest physically based guide to decide on the value of the threshold is in fact radial in nature rather than diametral.

## Computational details

### Clustering algorithms

The two algorithms, QTC^4^ and RTC,^2,3^ require as input the matrix of RMSDs between all pairs of configurations.

In its *m*^*th*^ iteration, the QTC algorithm scans the matrix in search for the configuration with the largest number of neighbors in order to generate the *m*^*th*^ cluster. Specifically, each configuration not clustered in the previous *m −* 1 iterations (we shall refer to them as the *available* configurations) will be considered both as seed of a tentative cluster and as potential neighbor of all other seeds. We use the term *tentative cluster* to refer to a seed and its neighbors before the neighborhood sizes of all seeds are compared to select the actual cluster. For each seed, its neighbors will be determined as follows: From all other available configurations, the one that upon its addition extends the least the diameter of the cluster, while fulfilling the condition that the diameter must remain smaller than the threshold, is taken as the next neighbor and included in the seed’s tentative cluster. This process is repeated until no remaining available configuration fulfills the threshold, at which point the tentative cluster for that seed is complete. Note that within an iteration all available configurations are tested as potential neighbors of each one of the available seeds. Once the tentative clusters for all available seeds have been obtained, the one with the largest number of elements is promoted to constitute the *m*^*th*^ cluster, and all elements of that cluster are removed from the pool of available configurations, thus finalizing the *m*^*th*^ iteration. The algorithm is stopped when there are no more configurations available or new clusters fall below a preset minimum number of elements.

The RTC algorithm differs from the QTC one in the way the neighbors of a seed are determined at each iteration: All available configurations at a distance from the seed smaller than the threshold are taken as elements of the seed’s tentative cluster. Thus, the RTC algorithm avoids the double loop per seed that characterizes the QTC algorithm —to find the next element of the tentative cluster (inner loop over available configurations) until no other configurations fulfilling the diameter threshold are available (outer loop).

The clusterings were performed with inhouse software reproducing exactly the algorithms described here. Results using established software implementations (available free of charge) on the tau-polypeptide example are provided as Supporting information (SI) for comparison. For the RTC case, results found in SI were obtained using the McLachlan algorithm^6^ as implemented in ProFit v3.3 (http://www.bioinf.org.uk/software/profit/) for the RMSD calculation and the RTC algorithm as implemented in HADDOCK v2.0^7^ (*cluster_struc*, https://www.bonvinlab.org/software/haddock2.2/) for the clustering. For the QTC case, results found in SI were obtained using the implementation published by González-Alemán et al. ^1^ (https://github.com/rglez/QT). The results are exactly the same, with small differences in the QTC case due to implementation details that are explained in the SI document and conform in both cases with the QTC algortihm.

### MD simulation data

The trajectory of the tau polypeptide was downloaded from https://github.com/LQCT/BitQT/blob/master/examples/aligned_original_tau_6K.dcd, together with the reference PDB file https://github.com/LQCT/BitQT/blob/master/examples/aligned_tau.pdb. It corresponds to the exact same trajectory used by González-Alemán et al. ^1^ in their comparison of the QTC and RTC clustering algorithms. The trajectory contains 6001 configurations of the polypeptide. We used the backbone N, H, C_*α*_, C, O atoms of residues Lys_2_ to Asp_11_, i.e., 50 atoms in total, for least-squares fitting and RMSD calculation. The two terminal residues, Gly_1_ and Leu_12_ and their capping groups were left out because they are relatively free to rotate and would introduce unnecessary noise in the clustering of the rest of the structure. Likewise, we excluded the side chains since one generally focuses on the backbone to define a fold and side chains would only introduce noise. This selection of atoms is clearly different from that used by González-Alemán et al. ^1^ (all atoms), but this is irrelevant for the questions addressed here.

To generate points for the example in two dimensions (not that this is important) we simply took the x and y coordinates of the backbone N atom of Lys_2_ after least-squares fitting of all configurations to configuration number 2910. A subset of 1501 elements was then constructed by selecting 1 element every 4, starting with element 1.

## Results and discussion

### Theoretical framework and properties

We note that throughout this article we use the term *diameter* in its generalized form, i.e., as the largest distance between any two points on the boundary of a closed geometric figure (in this case a cluster). Likewise, we use the term *sphere* as short-hand for (*n −* 1)-sphere, defined as the (*n −* 1)-dimensional boundary of an (*n*-dimensional) *n*-ball.

Let *S*_*m*_ = {**x**_*i*_ *≡* (*x*_*i*,1_, …, *x*_*i,n*_) ∈ ℝ^*n*^ : *i* ∈ *I*_*m*_} be a set of Euclidean vectors in a Cartesian frame (for convenience, we shall also refer to **x**_*i*_ as a *point* in that frame) representing the *N*_*m*_ configurations of the molecule that are available for clustering at the *m*^*th*^ iteration of the RTC or QTC algorithm, where *n* is the number of coordinates that will be used for the RMSD calculation, *I*_*m*_ = {*i* ∈ ℕ : 1 ≤ *i* ≤ *N*_1_, *i* ∉ *J*_*m*_} is the set of indices of the elements of *S*_1_ that are available for clustering at the *m*^*th*^ iteration and *J*_*m*_, with *J*_1_ = ∅ and *J*_*m>*1_ = {*j* ∈ ℕ : 1 ≤ *j* ≤ *N*_1_, **x**_*j*_ ∈ *C*_*l*_, 1 ≤ *l* ≤ *m −* 1}, is the set of indices of the elements of *S*_1_ that have been already clustered in previous iterations, where *C*_*l*_ stands for the cluster set defined in iteration *l* (see below).

Let **x**_*k*_ ∈ *S*_*m*_ be the seed for a tentative cluster of elements of *S*_*m*_ and *A*_*m,k*_(*θ*) = {**x**_*i*_ ∈ *S*_*m*_ : *RMSD*_*ki*_ *< θ*} the set of elements within a RMSD-threshold *θ* ∈ ℝ_*>*0_ from **x**_*k*_. Note that *RMSD*_*ki*_ = ‖ **x**_*i*_ *−* **x**_*k*_ ‖ (*n*_*a*_)^*−*1*/*2^, where *n*_*a*_ is the number of atoms involved in the RMSD calculation (in principle, *n* = 3 *× n*_*a*_). Then, we define *A*_*m*_(*θ*) = {*A*_*m,k*_(*θ*) : *k* ∈ *I*_*m*_} as the collection of such sets for all available seeds and *B*_*m*_(*θ*) = {|*A*_*m,k*_(*θ*)| : *k* ∈ *I*_*m*_}, where |*A*_*m,k*_(*θ*)| stands for the cardinality of *A*_*m,k*_(*θ*), as the collection of corresponding set sizes.

In the RTC algorithm *A*_*m,k*_(*θ*) is the tentative cluster “proposed” by seed **x**_*k*_, which shall be then compared to the tentative clusters “proposed” by all other seeds. Thus, we define *D*_*m*_(*θ*) = {*A*_*m,k*_(*θ*) ∈ *A*_*m*_(*θ*) : |*A*_*m,k*_(*θ*)| = max (*B*_*m*_(*θ*))} as the collection of sets with the largest number of elements and *E*_*m*_(*θ*) = {*k* ∈ *I*_*m*_ : *A*_*m,k*_(*θ*) ∈ *D*_*m*_(*θ*)} as the collection of corresponding indices. The *m*^*th*^ cluster (output of the *m*^*th*^ iteration of the algorithm) is then defined as *C*_*m*_(*θ*) = {*A*_*m,k*_(*θ*) ∈ *D*_*m*_(*θ*) : *k* = *f* (*E*_*m*_(*θ*))}, where *f* is a function that returns one element from a set, typically the function min(), in which case *C*_*m*_(*θ*) would be the set with lowest index from those with largest number of elements.

To impose the condition that the diameter of *C*_*m*_(*θ*) be smaller than *θ*, as done in the QTC algorithm, we need to define a new set *F*_*m,k*_(*θ*) = *p*(*A*_*m,k*_(*θ*)), where *p* is an element-selection procedure, i.e., *F*_*m,k*_(*θ*) ⊂ *A*_*m,k*_(*θ*), such that *RMSD*_*ij*_ *< θ*, ∀ **x**_*i*_, **x**_*j*_ ∈ *F*_*m,k*_(*θ*). *F*_*m,k*_(*θ*) is, in this algorithmic context, the tentative cluster “proposed” by seed **x**_*k*_, which shall be compared to the tentative clusters “proposed” by all other seeds. Thus, as done for the tentative clusters in the RTC case, we define *F*_*m*_(*θ*) = {*F*_*m,k*_(*θ*) : *k* ∈ *I*_*m*_} as the collection of such sets for all available seeds and redefine *B*_*m*_(*θ*) = {|*F*_*m,k*_(*θ*)| : *k* ∈ *I*_*m*_} as the collection of corresponding set sizes. Accordingly, we redefine *D*_*m*_(*θ*) = {*F*_*m,k*_(*θ*) ∈ *F*_*m*_(*θ*) : |*F*_*m,k*_(*θ*)| = max (*B*_*m*_(*θ*))} and *E*_*m*_(*θ*) = {*k* ∈ *I*_*m*_ : *F*_*m,k*_(*θ*) ∈ *D*_*m*_(*θ*)}. The *m*^*th*^ cluster is then defined as *C*_*m*_(*θ*) = {*F*_*m,k*_(*θ*) ∈ *D*_*m*_(*θ*) : *k* = *f* (*E*_*m*_(*θ*))}.

Note that the latter cluster definition is not specific for the QTC algorithm, but general for a group of diameter-based algorithms. This is because, as it is defined, the procedure *p* is not unique. That is, the condition *RMSD*_*ij*_ *< θ*, ∀ **x**_*i*_, **x**_*j*_ ∈ *F*_*m,k*_(*θ*), *F*_*m,k*_(*θ*) ⊂ *A*_*m,k*_(*θ*) can be satisfied by different selection procedures *p*, leading to different subsets *F*_*m,k*_(*θ*) of *A*_*m,k*_(*θ*). For example, the tentative cluster *F*_*m,k*_(*θ*) can be grown starting from its seed following the procedure by Heyer et al. ^4^ and described under Computational details (the procedure *p* implemented in the QTC algorithm), or could be grown following a specific sequence of configurations (e.g., time sequence), i.e., testing at each step if the next configuration in sequence satisfies the diametral threshold instead of searching for the configuration that minimizes the increase in cluster diameter (note that this would produce different *F*_*m,k*_(*θ*) subsets for different configuration sequences). Another variant of *p* could be searching for the subset *F*_*m,k*_(*θ*) with the largest number of elements, etc.

While the RTC and QTC algorithms are generally presented as invariant to permutation (referring to the order in which the algorithm evaluates the seeds or the order in which the configurations are tested for inclusion in a seed’s tentative cluster), they are strictly not. This is because the collection *D*_*m*_(*θ*) defined above may contain more than one set. Indeed, when using these algorithms in practice, at any given iteration it is relatively common to see two or more seeds tie as the ones generating the tentative clusters with the largest number of elements (this is precisely why *f* is needed in the definition of *C*_*m*_(*θ*)), in which case both algorithms become configuration-order sensitive (for a given function *f*). It is true, however, that ties tend to occur between seeds that are very close in space, thus having a relatively small impact on the clustering.

We shall use the term *cluster shape* to refer to the convex hull of the set of points that belong to the cluster. Thus, the cluster shape with maximum volume is for either algorithm a sphere, with radius *θ* for RTC clusters and *θ/*2 for QTC ones. Note that we can define three types of geometric centers for both RTC and QTC clusters. The first type is the center of the neighbor-search volume (a sphere) and corresponds to the position of the seed (as we have seen above, both algorithms search within *A*_*m,k*_(*θ*)). For this reason, it is customary to refer to the seed, particularly in the RTC case, as the central element of the cluster, but this is, as we shall see, misleading if we give it a spatial significance. The second type is the geometric center of the cluster shape, which will only coincide with the first one if the cluster is spherical (RTC case) or spherical and centered on the seed (QTC case). And the third type is the geometric center of the cluster elements, which will only coincide with the second one if the spatial distribution of points in the cluster is homogeneous (which is rare). Thus, it is in fact common for the seeds of even RTC clusters to be relatively distant from the geometric center of the cluster shape and/or the geometric center of the cluster elements.

The seed of a QTC cluster tends in fact to be close to the cluster’s boundary. This is due to the specific procedure *p* that selects the elements of *F*_*m,k*_(*θ*) from *A*_*m,k*_(*θ*). As described under Computational details, at each step in the process of recruiting new elements for the tentative cluster *F*_*m,k*_(*θ*), the point that minimizes the increase in diameter while fulfilling the diametral threshold *θ* is selected. Thus, the first steps of the procedure are highly determined by the local distribution of points around the seed, which is never equal in all directions. This will introduce an early bias or dominant direction and sense for the cluster’s growth, which may be more or less prominent depending on the exact spatial distribution of points. In cases in which the distribution induces a marked directionality (which could be relatively frequent as we shall see in the 2D example below), and assuming a spherical cluster, the cluster will tend to grow in eccentric spherical layers away from the seed, leaving the seed at or very close to the cluster’s boundary. Note that it is the local distribution of points around the seed that primarily determines the direction and sense of growth of *F*_*m,k*_(*θ*), rather than the global spatial distribution of points in *A*_*m,k*_(*θ*). Therefore, *F*_*m,k*_(*θ*) is not necessarily the subset of *A*_*m,k*_(*θ*) with highest cardinality. Another consequence of this is that, unlike for RTC clusters, for QTC clusters the relation |*C*_*m*_(*θ*)| *>* |*C*_*m*+1_(*θ*)| does not need to hold: the direction of growth of the tentative cluster around a given seed **x**_*k*_ may in some cases be less optimal for set *S*_*m*_ than for set *S*_*m*+1_, so that we may have |*F*_*m,k*_(*θ*)| *<* |*F*_*m*+1,*k*_(*θ*)|, which can eventually lead to a situation in which |*F*_*m,k*_(*θ*)| *<* |*F*_*m,j*_(*θ*)|, where *F*_*m,j*_(*θ*) = *C*_*m*_(*θ*), and |*F*_*m*+1,*k*_(*θ*)| *>* |*F*_*m,j*_(*θ*)|, where *F*_*m*+1,*k*_(*θ*) = *C*_*m*+1_(*θ*). In practice, however, the fact that *C*_*m*_(*θ*) is selected from a collection *F*_*m*_(*θ*) of overlapping tentative clusters *F*_*m,k*_(*θ*), makes this potential inversion of cluster-size order very infrequent, and we have only observed it in two cases in the examples below, for very small (irrelevant) clusters.

To adequately compare the RTC and QTC algorithms in practical applications, one needs to keep in mind that *θ* is a radial threshold (i.e., *θ*_*r*_) in the RTC algorithm and a diametral threshold (i.e., *θ*_*d*_) in the QTC algorithm. If the spatial distribution of points is such that the diameters of the RTC clusters are approximately equal to two times the threshold *θ*_*r*_ in at least one direction, the extreme case being spherical clusters, one should use *θ*_*r*_ = *θ/*2 as RTC threshold and *θ*_*d*_ = *θ* as QTC threshold for comparable results. However, as we shall see in the tau-polypeptide example, the distribution of points has rarely these characteristics when working with *n*-dimensional data representing molecular configurations from computer simulation. First, the region of this *n*-dimensional space corresponding to a conformer of the molecule (where the term *conformer* may be taken as a very well defined structure or a broader structural state, depending on the purpose of the clustering) will generally have an irregular shape and its immediate surrounding may be void in many directions, so that even if the algorithm tries to mix in points corresponding to neighbor conformers (when the threshold is too big) in many spatial directions there might be simply nothing to mix in. Second, the density of points in this *n*-dimensional space is typically reduced in clustering exercises due to the selection of only one structure every so many simulation steps, which has the effect of blurring the underlying cluster topology (if such should physically exist). Therefore, in order to have equally sized clusters, the relation between the RMSD thresholds *θ*_*r*_ (RTC) and *θ*_*d*_ (QTC) will be in practice *θ*_*d*_*/*2 ≤ *θ*_*r*_ *< θ*_*d*_.

A more important question, however, is how to choose the threshold. Typically, the option of choice in the literature is trial and error: both algorithms, particularly RTC, are sufficiently fast that one can actually run them several times with different *θ* values, until the outcome satisfies any chosen criteria. The criterion is often visual, i.e., the structures in a cluster “look the same”.^1^ While this may fit the purpose in some types of studies, it is actually a weak criterion from a physical stand point. As a general rule, before clustering data points in an *n*-dimensional space, one may want to query this space to gather information on the distribution of data points in it. This is also what makes physical sense in this case, since the space in question is the molecule’s configuration space (reduced to the number of coordinates used for RMSD calculation and the given ensemble sample). As will be shown for the tau-polypeptide example below, the simplest effective way to query this space is by calculating the distribution of RMSD values. This can be done for the full (half) RMSD matrix, but the mixing of underlying distributions generally reduces its informative value. Instead, we propose that a first tentative clustering may be performed to calculate the RMSD distribution for each of the seed elements of the most populated clusters (as representatives of the high-density regions of interest), i.e., taking the respective full columns (or rows) from the RMSD matrix. In many cases, these distributions will show a first region of relatively high probability density, followed by a deep and then a second, larger increase in probability density. This deep in the probability density corresponds to the end of the first layer of configurations around the seed, and may therefore be interpreted as a (spherical) region of conformational transition. After comparing the distributions for the seed elements of the main clusters, this information can be used to assess the threshold for a second, final clustering. Note that this information is radial in nature, and it therefore leads to a value for *θ*_*r*_ rather than *θ*_*d*_.

In the next subsections we illustrate some of the properties discussed above of the RTC and QTC algorithms with two examples.

### Example case in 2D

Figure 1 shows the clustering of the set of 1501 points in two dimensions (see Computational details) performed with the RTC (panel A) and QTC (panels B, C) algorithms. This is a good example of a major limitation of fixed-size clustering algorithms: If the data has no particular underlying structure or the threshold is completely inadequate, these algorithms will still partition the data according to the chosen threshold. The question of how to choose a threshold is therefore of particular significance, even if we will ignore it in this first example. Note that, as mentioned under Theoretical framework and properties, for densely populated spaces with few void regions a selection of thresholds such that *θ*_*r*_ = *θ*_*d*_*/*2, where *θ*_*r*_ is the threshold used with the RTC algorithm and *θ*_*d*_ is the threshold used with the QTC algorithm, produces equally sized clusters in the two cases. As also mentioned, while the seeds of the RTC clusters tend to be closer to the centroids of their cluster shapes, the seeds of the QTC clusters tend to be closer to the clusters’ boundaries.

**Figure 1:**
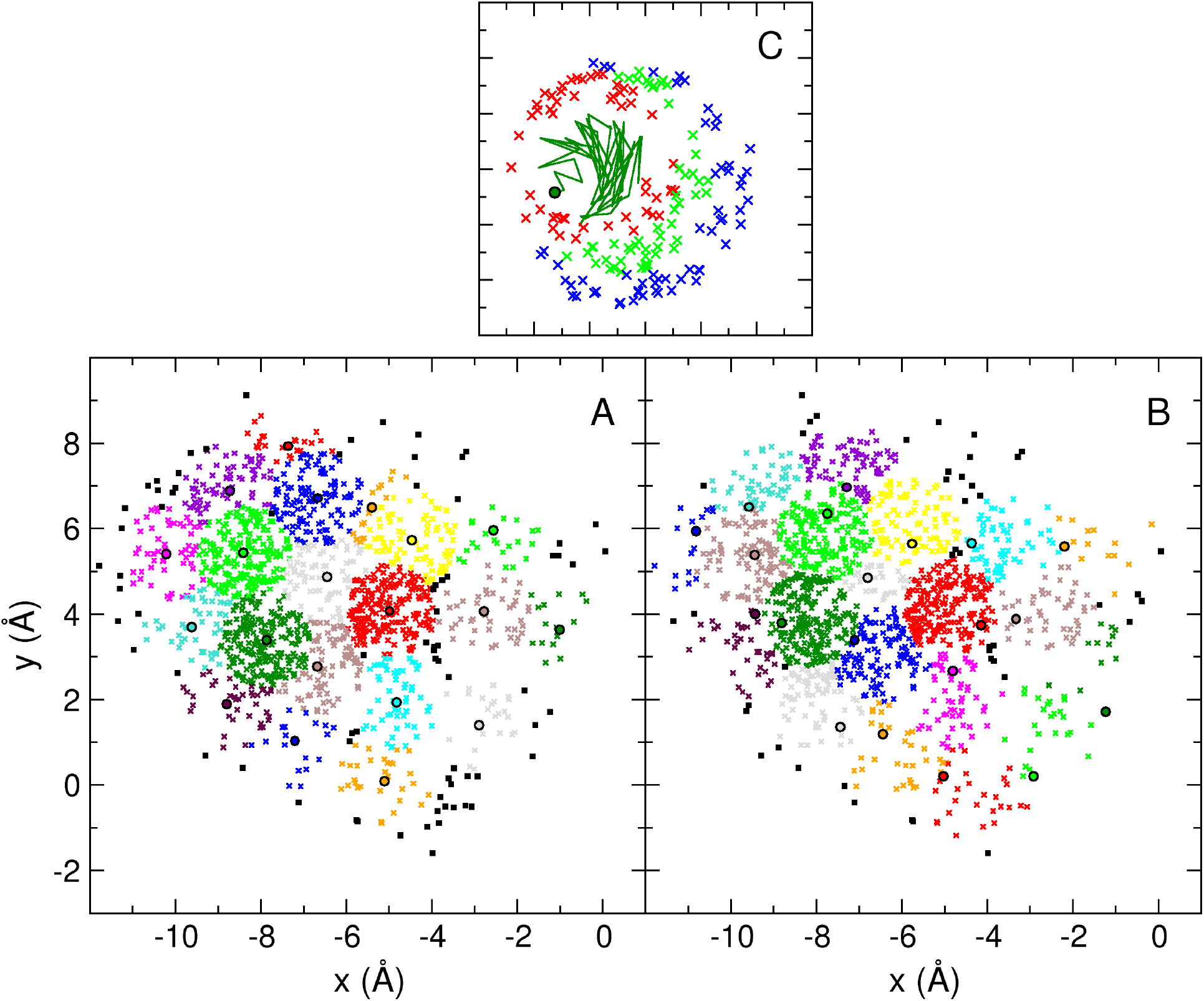
Representation of the first 20 clusters of 1501 points in 2D. **A**. RTC algorithm, *θ* = 1.1 Å. **B**. QTC algorithm, *θ* = 2.2 Å. The color sequence is the same in both panels, starting with dark green for the first cluster. The seed elements of the clusters are indicated with a black circle (with area matching the cluster color). Points belonging to clusters with index higher than 20 are indicated as black squares. **C**. Detail of the process of generation of cluster 1 from panel B (dark green): Initial growth from the seed (dark-green circle) is indicated as a dark-green trajectory. Elements in red, light green and blue, correspond to successive phases in the growth of the cluster, in this order.

Panel C illustrates the generation process of QTC clusters, using the first cluster from panel B (dark green) as example. The initial steps are shown as a trajectory starting from the seed (dark-green circle). Consecutive phases in the growing of the cluster are illustrated with elements in red, light green and blue, in this order. The directionality of the growth, away from the seed in eccentric circular layers, can be clearly observed.

### Clustering of structures from a MD trajectory

Following the strategy described above to infer a physically meaningful threshold, we first used the RTC algorithm to perform a clustering of the 6001 configurations of the taupolypeptide, in order to identify points in the 150-dimensional space (50 atoms *×* 3 coordinates) that are in regions of high density. In this case we chose the seed elements of the six most populated clusters to then examine the distribution of RMSDs between each of these elements and the other 6000 configurations. In order to see how the distributions may vary depending on the threshold used for this initial clustering, we chose six evenly spaced thresholds between 0.12 and 0.17 nm. The results are shown in Figure 2.

**Figure 2:**
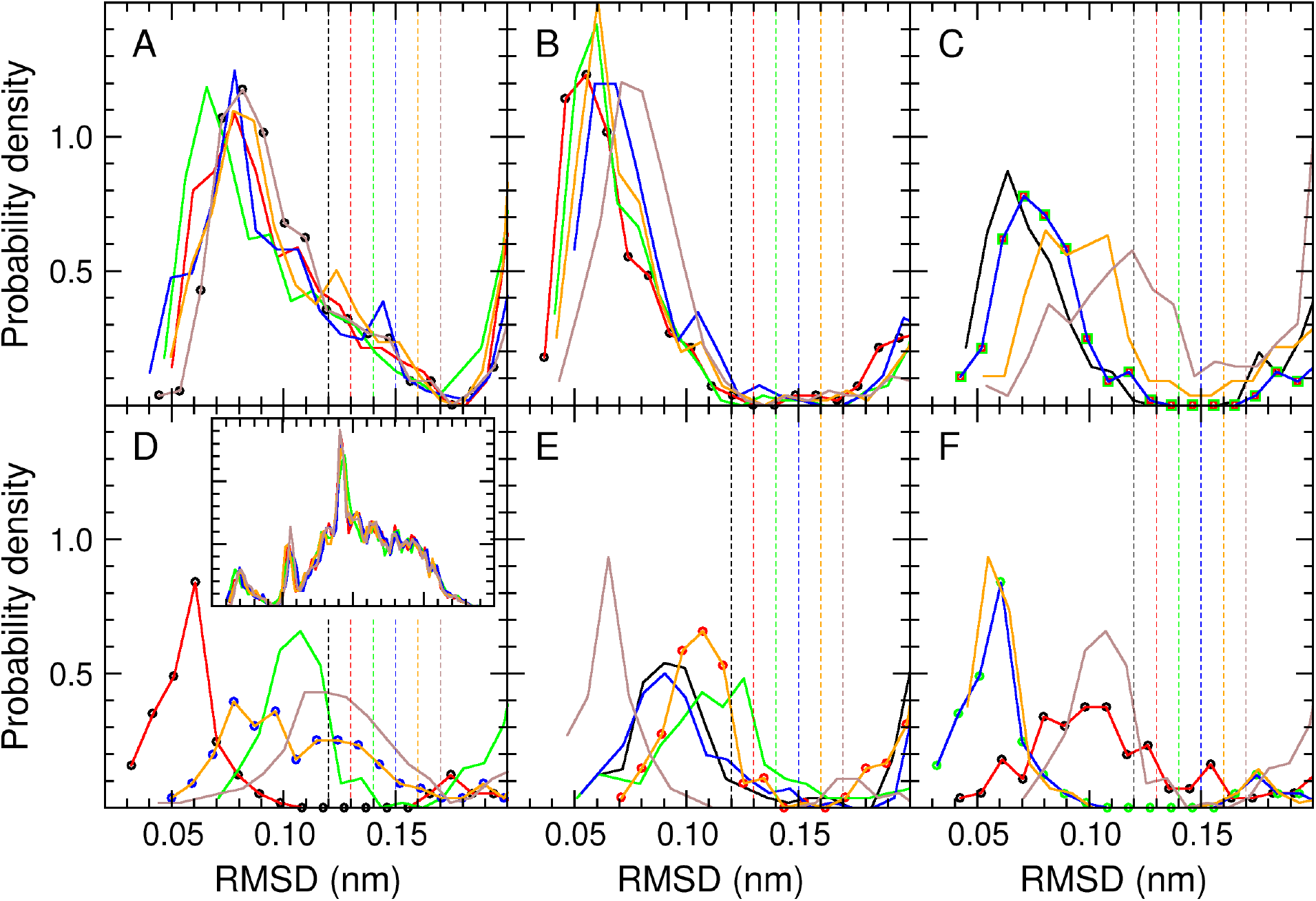
RMSD distributions for the seeds of the six most populated clusters, after RTC clustering with six different thresholds. Each curve corresponds to the distribution of the RMSDs between the given seed and all other 6000 configurations. The distributions are cut at 0.20 nm for clarity (the inset in panel D shows the full distributions from panel A as example). Panels **A** to **F** correspond to the distributions for the seeds of clusters 1 to 6, respectively. Each panel contains the distributions for six seeds, resulting from RTC clusterings with *θ*_*r*_ values of 0.12 nm (black), 0.13 nm (red), 0.14 nm (green), 0.15 nm (blue), 0.16 nm (orange) and 0.17 nm (brown). The dashed vertical lines show the positions of the thresholds (with colors matching the corresponding distributions). When two distributions overlap (i.e., clusterings with different thresholds produce the same seeds for the given cluster number), the overlapping curves are replaced by circles of the corresponding color.

In all distributions, an initial region corresponding to a first layer of configurations around the seed can be distinguished, after which the probability density goes down to zero or close to it. The distributions for the seeds of the first, most populated clusters tend to be less sensitive to the threshold used for the clustering, as expected. This is not so much because the clusters are more populated, but because they are generated first, i.e., the following clusters are affected by which configurations have or have not been already taken by the previous ones. Although the RMSD value at which the probability density reaches zero differs for the different distributions, 0.17 nm stands out as a possible consensus threshold: it is a point at which the probability density either reaches zero (notably for the seeds of cluster 1) or has not yet recovered significantly from zero.

Based on these observations we focused on the clusterings performed with *θ*_*r*_ values of 0.12 nm, i.e., the lowest value that seems adequate for some of the distributions shown in Figure 2, and 0.17 nm. To define corresponding *θ*_*d*_ thresholds for the QTC algorithm we looked at the diameter of cluster 1 in each of these two RTC clusterings. Note that cluster 1 is our best guide because its diameter is not conditioned by previous clusters. The diameter was 0.199 nm for 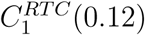, and 0.278 nm for 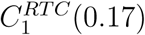. With these reference values and taking into account that the probability density for a RMSD distance above 0.25 nm in 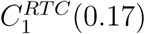 is very small (only 17 elements at distances between 0.25 and 0.278 nm), we decided to choose *θ*_*d*_ values of 0.20 nm and 0.25 nm for the QTC algorithm.

Note that, as already discussed, the diameter of an RTC cluster should be generally expected to be smaller than 2*θ*_*r*_ for actual data sets from simulation, as confirmed by the values indicated in the previous paragraph. The reason for this is illustrated in Figure 3. This figure shows, for every pair of elements (excluding the seed) of clusters 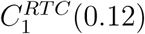 (panel A) and 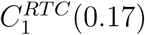 (panel B), the relation between the sum of their RMSD distances to the seed and the angle between the vectors associated to these distances. Thus, while there are pairs of points for which the sum of RMSDs adds up indeed to 2*θ*_*r*_, i.e., 0.24 nm (panel A) and 0.34 nm (panel B), the angle between the corresponding vectors is in no case close to 180^*o*^, which precludes the maximum diameter from being reached. This figure also shows that, as expected (see panel A in Figure 2), the point density is abruptly cut (solid wall at 0.24 nm) when using the 0.12 nm threshold (panel A), while a threshold of 0.17 nm leads to a well-defined cluster (panel B).

**Figure 3:**
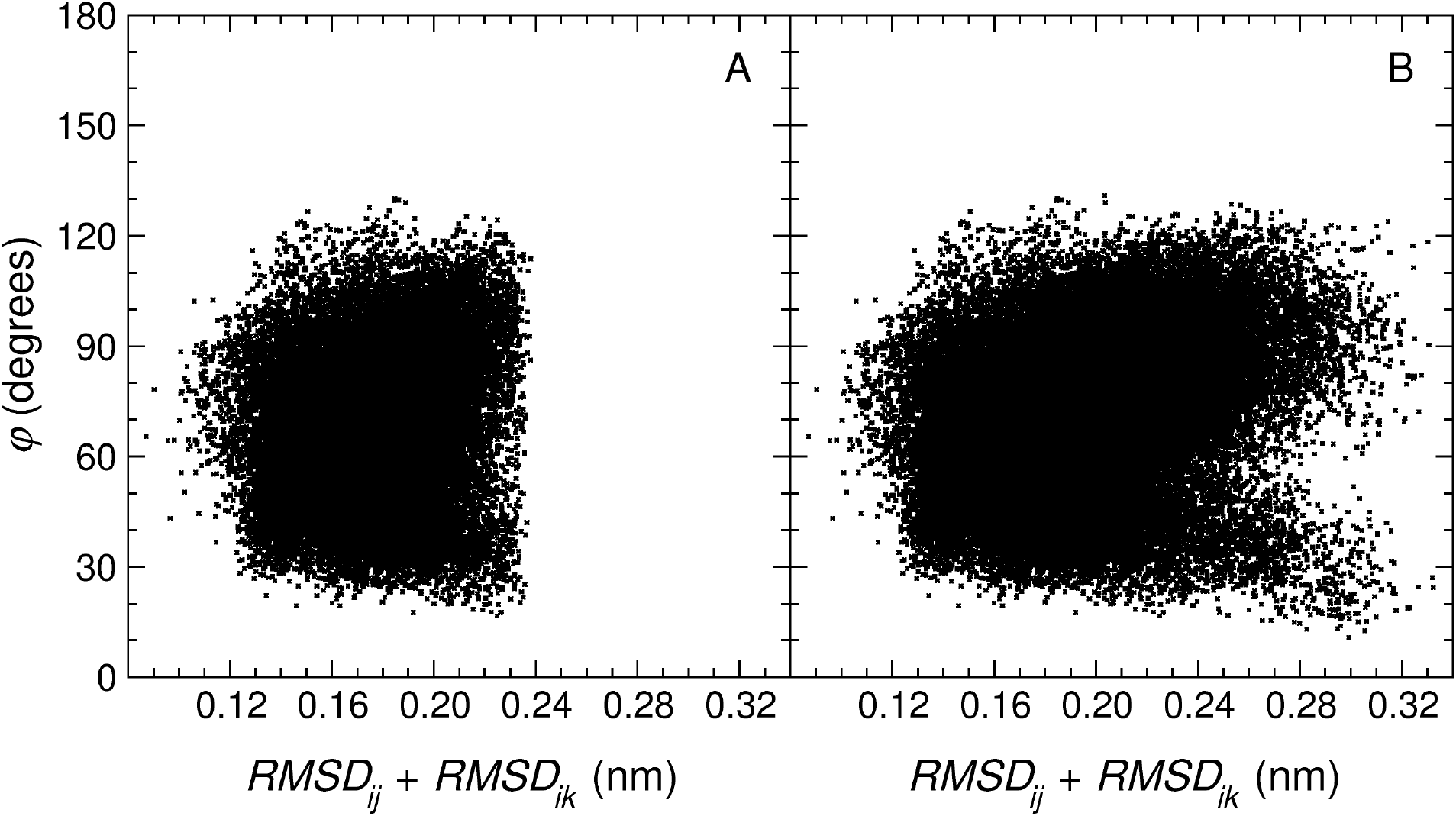
Relation between distance to the seed and corresponding angle for every pair of elements of a cluster (excluding the seed). Specifically, *RMSD*_*ij*_ + *RMSD*_*ik*_, for all **x**_*j*_, **x**_*k*_ ∈ *C*_1_(*θ*_*r*_), *j, k* ≠ *i*, where **x**_*i*_ is the seed of *C*_1_(*θ*_*r*_) and *θ*_*r*_ has the values 0.12 nm (**A**) and 0.17 nm (**B**), against the angle *ϕ* between the vectors **x**_*ij*_ and **x**_*ik*_, where **x**_*ij*_ = **x**_*j*_ *−* **x**_*i*_.

Figure 4 shows the distribution of RMSD values within each of the first ten clusters, for the RTC clustering with *θ*_*r*_ = 0.12 nm and the QTC clustering with *θ*_*d*_ = 0.20 nm. Seven of the clusters, including the first four, overlap almost perfectly. For three of the clusters, the QTC algorithm appears to have a higher tendency to populate the far-right side of the distribution.

**Figure 4:**
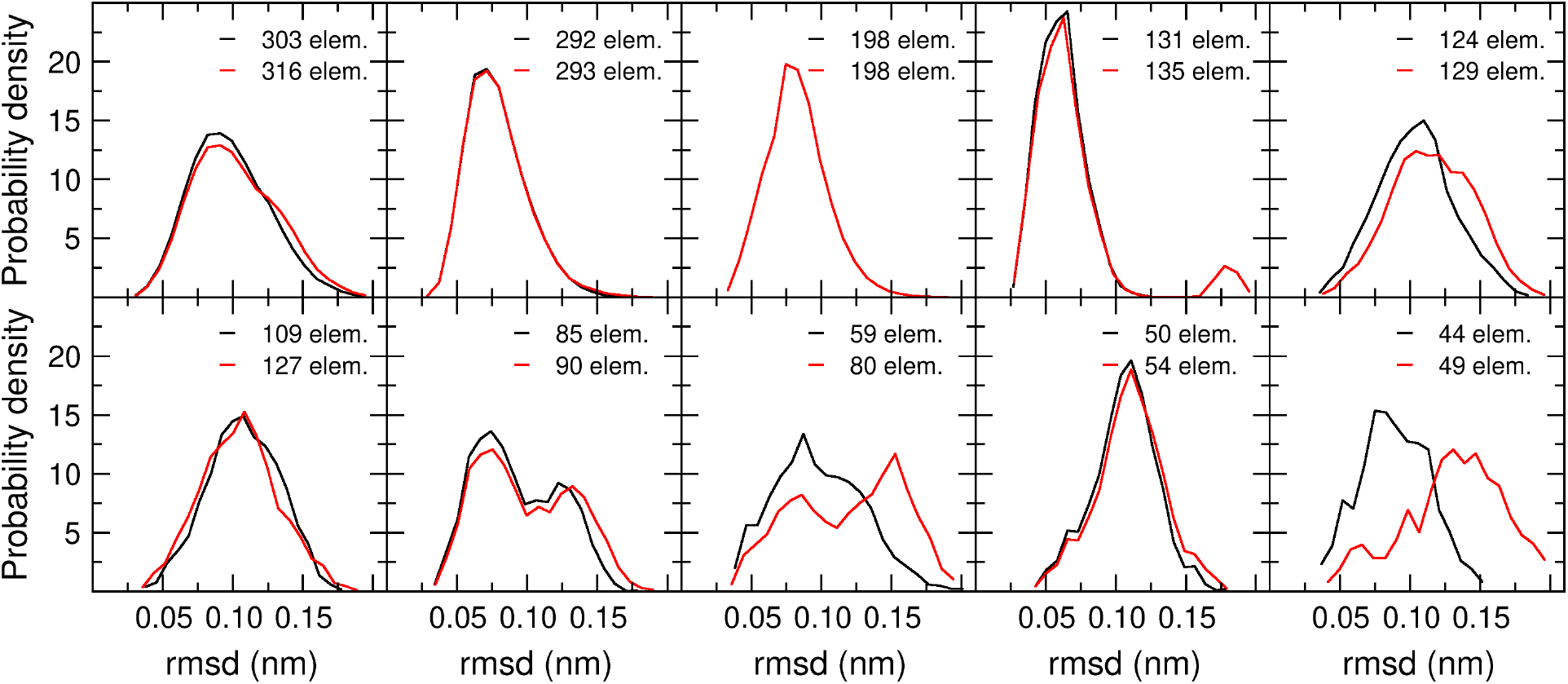
RMSD distributions for the first ten clusters, using the RTC algorithm with *θ*_*r*_ = 0.12 nm (black) and the QTC algorithm with *θ*_*d*_ = 0.20 nm (red). The number of elements in the cluster is for each case indicated.

Figure 5 shows the corresponding distributions for the RTC clustering with *θ*_*r*_ = 0.17 nm and the QTC clustering with *θ*_*d*_ = 0.25 nm. It becomes here more apparent that the QTC algorithm has a higher tendency to generate split distributions and populate the far-right side of the distribution. For example, it can be observed that an artificial cluster 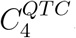, containing two different populations with similar weights, has displaced by one position in the ranking the QTC clusters that correspond to 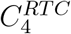 and 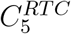.

**Figure 5:**
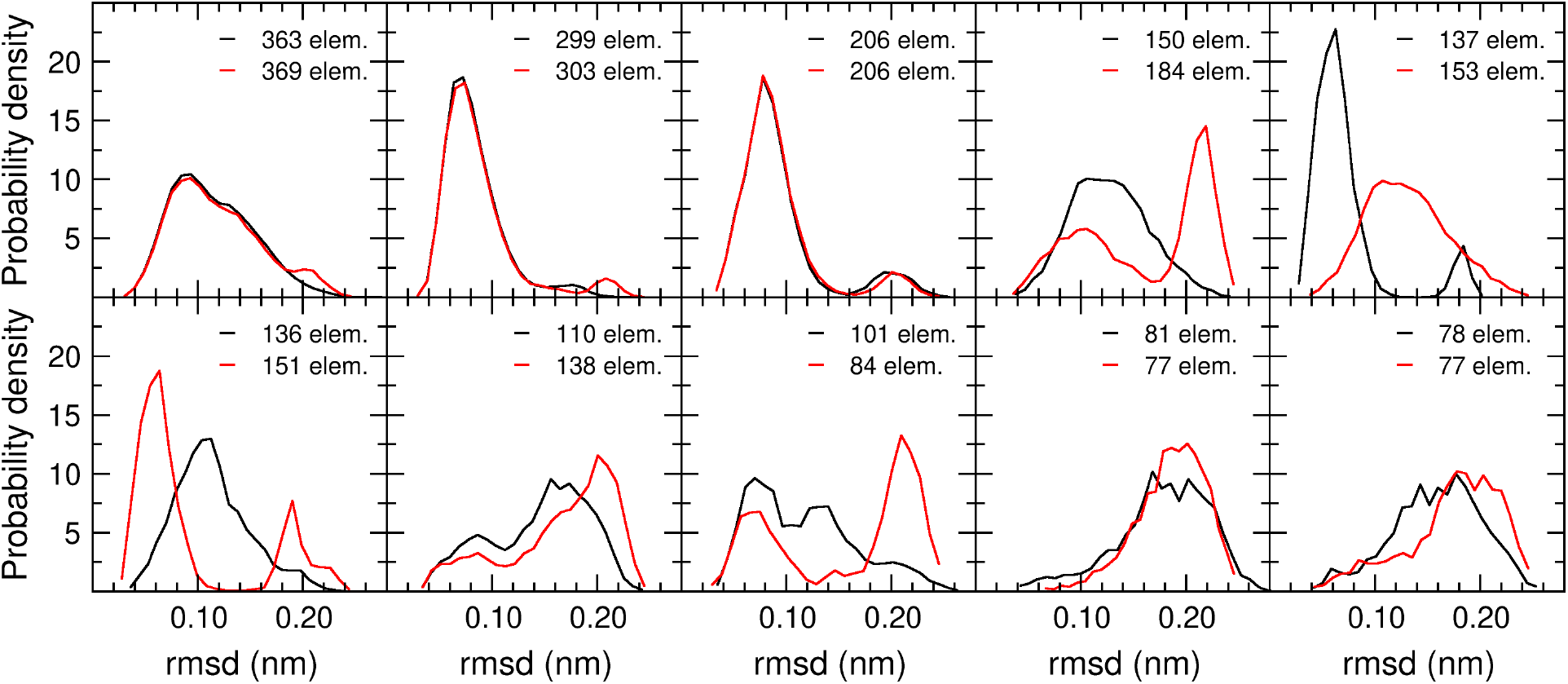
RMSD distributions for the first ten clusters, using the RTC algorithm with *θ*_*r*_ = 0.17 nm (black) and the QTC algorithm with *θ*_*d*_ = 0.25 nm (red). The number of elements in the cluster is for each case indicated.

We obtained for each cluster *C*_*m*_ shown in Figures 4 and 5 the center of geometry of the elements of the cluster, 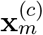, and then calculated the RMSD between the seed element and this point, as well as the radius of gyration of the cluster. Note that the radius of gyration of the *m*^*th*^ cluster is here defined as:

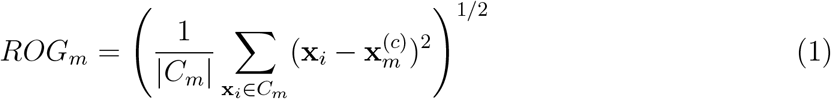

The results are shown in Figure 6. As anticipated, panel A confirms that while the seeds of QTC clusters are in a majority of cases further from the center of geometry of the cluster elements than the seeds of RTC clusters, the latter are clearly off-center also. We therefore suggest to avoid, in either case, the term *central element* to refer to the seed element. Panel B illustrates that, except for cases in which a tight match between the RMSD distributions in Figures 4 and 5 exists, the RTC clusters tend to be more compact (lower ROG) than the QTC clusters.

**Figure 6:**
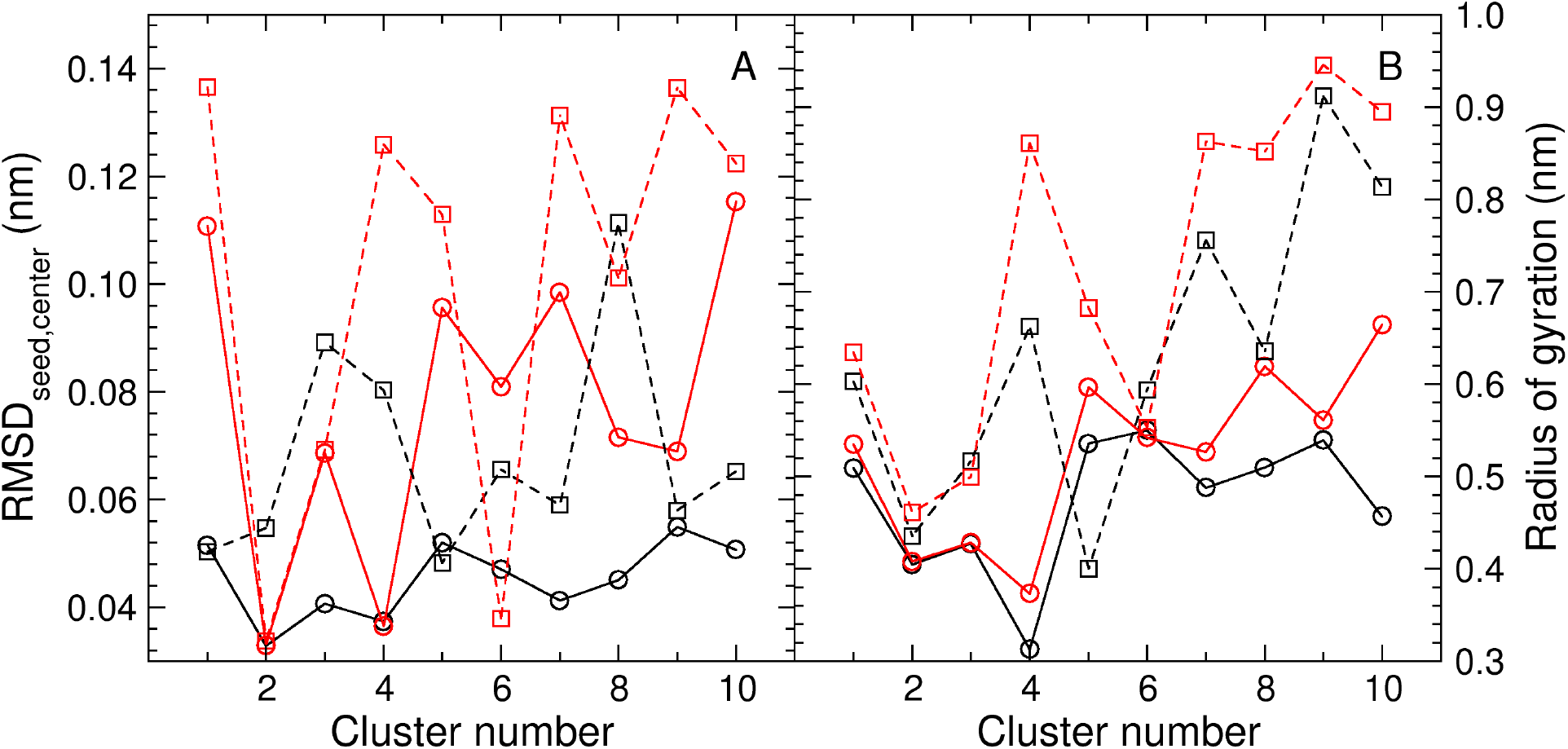
Distance of seed from the center and cluster compactness. **A**. RMSD between the seed element and the center of geometry of the elements in the cluster. **B**. Radius of gyration of the cluster. Black circles (black solid line): RTC clustering with *θ*_*r*_ = 0.12 nm. Red circles (red solid line): QTC clustering with *θ*_*d*_ = 0.20 nm. Black squares (black dashed line): RTC clustering with *θ*_*r*_ = 0.17 nm. Red squares (red dashed line): QTC clustering with *θ*_*d*_ = 0.25 nm.

When looking at the total number of clusters generated by the two algorithms we see that they differ remarkably (see Supporting information). Thus, while the number of clusters generated by the RTC algorithm using *θ*_*r*_ values of 0.12 nm and 0.17 nm is 1338 and 493, respectively, corresponding numbers for the QTC algorithm with *θ*_*d*_ values of 0.20 nm and 0.25 nm are 599 and 276, respectively. However, focusing on the upper part of the ranking list, we see that for RTC with *θ*_*r*_ = 0.12 nm the number of clusters with 100 or more elements is 6 and these cover 19% of the overall population, while for QTC with *θ*_*d*_ = 0.20 nm the number of clusters is also 6 and they cover 20% of the population. If we extend this to clusters with 10 or more elements, the numbers start to diverge, with RTC producing 98 clusters that cover 50% of the population and QTC producing 161 clusters covering 70% of the population. The trends are similar but the results less divergent for the comparison between RTC with *θ*_*r*_ = 0.17 nm and QTC with *θ*_*d*_ = 0.25 nm. Here, the RTC algorithm generates 8 clusters with 100 or more elements covering 25% of the population and 149 clusters with 10 or more elements covering 82% of the population, while the QTC algorithm generates 7 clusters with 100 or more elements covering 25% of the population and 150 clusters with 10 or more elements covering 91% of the population. Thus, while the upper part of the ranking looks very much the same with the two algorithms, RTC produces many more small clusters at the end of the ranking.

How can we explain such large differences in the total number of clusters? The neighbor-search volume for the RTC algorithm is strictly spherical, while for the QTC algorithm it is a volume within a sphere, with the diameter as only shape restraint. This makes the QTC algorithm more flexible in terms of cluster shapes, allowing it to capture more of the elements that would lay just outside a cluster boundary in the RTC case. Thus, the RTC algorithm tends to generate many more artificial clusters with orphan elements in the lower part of the cluster ranking. On the other hand, the greater shape flexibility of the QTC algorithm makes it have a higher propensity to incorporate points from non-self neighbor densities in a cluster, thus mixing in it different populations. However, and as a conclusion, these differences between the two algorithms tend to be rather irrelevant in practice.

## Data and Software Availability

The data used in this study was downloaded from https://github.com/LQCT/BitQT/blob/master/examples/ as indicated under Computational details. Although the calculations shown here were performed with inhouse software, this calculations can be performed with a variety of software packages implementing the RTC and QTC algorithms (see some of the potential choices in González-Alemán et al. ^1^). We provide in Supporting Information (SI) a comparison between our results for the tau polypeptide and results using one of the alternative software options in each case. The results for the two software choices are exactly the same, with small differences in the QTC case due to implementation details that are explained in the SI document and conform in both cases with the QTC algortihm.

## Acknowledgement

The authors thank Joan-Emma Shea and Zachary Levine for permision to use their taupolypeptide simulation trajectories. This work received financial support from the Spanish Ministry for Science, Innovation and Universities (grant PID2019-111364RB-I00).

## Supporting Information Available

The following files are available free of charge.

- Tau_results_diff_software.xlsx: listing of clusters, including seed element and number of elements, from the tau-polypeptide example, with *θ*_*r*_ values of 0.12 nm and 0.17 nm (RTC clustering) and *θ*_*d*_ values of 0.20 nm and 0.25 nm (QTC clustering), corresponding to the data presented here (inhouse software) and computed with alternative available software, as indicated under Computational details.

## TOC Graphic

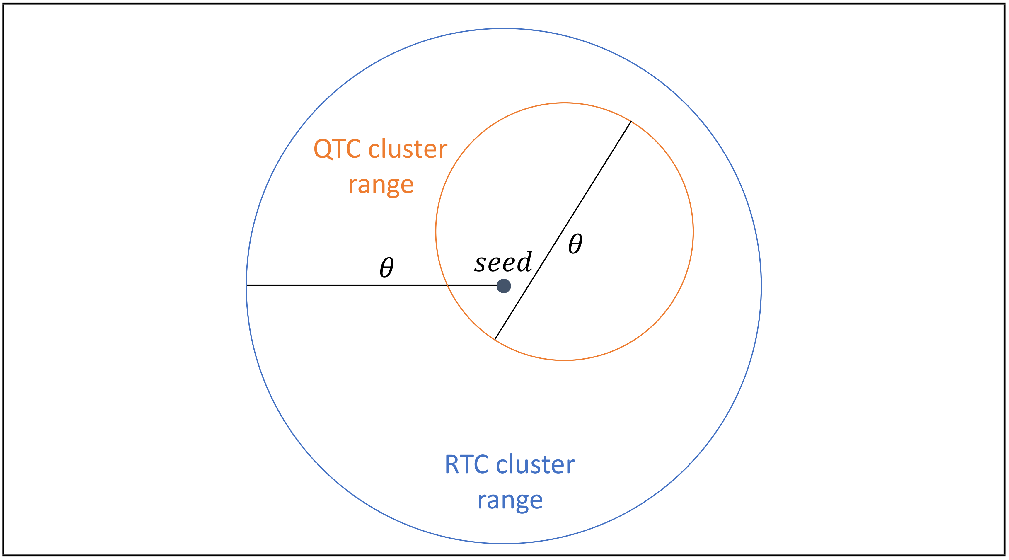

